# Restoration of Eastern oyster populations with positive density dependence

**DOI:** 10.1101/169540

**Authors:** Jacob L Moore, Brandon Puckett, Sebastian J Schreiber

## Abstract

Positive density dependence can create a threshold of population states below which extinction of the population occurs. The existence of this threshold, which can often be a complex, multi-dimensional surface, rather than a single point, is of particular importance in degraded populations for which there is a desire for successful restoration. Here, we incorporated positive density dependence into a closed, size- and age-structured integral projection model parameterized with empirical data from an Eastern oyster, *Crassostrea virginica*, population in Pamlico Sound, North Carolina. To understand the properties of the threshold surface, and implications for restoration, we introduced a general method based on a linearization of the threshold surface at its unique, unstable equilibrium. We estimated the number of oysters of a particular age (i.e. stock enhancement), or the surface area of hard substrate required (i.e. habitat enhancement), to move a population from an extinction trajectory to a persistent trajectory. The location of the threshold surface was strongly affected by changes in the amount of local larval retention. Traditional stock enhancement with oysters less than a year old (i.e. spat) required three times as many oysters relative to stock enhancement with oysters between ages three and seven, while the success of habitat enhancement depended upon the initial size distribution of the population. The methodology described here demonstrates the importance of considering positive density dependence in oyster populations, and also provides insights into effective management and restoration strategies when dealing with a high dimensional threshold separating extinction and persistence.

## INTRODUCTION

Many natural populations exhibit positive density dependence, or Allee effects, in which an increase in population size leads to an increase in per capita growth rate, or other components of fitness (Allee, 1931, 1949; Stephens et al., 1999; Courchamp et al., 1999). These Allee effects arise through various mechanisms, such at mate limitation or predator saturation (Courchamp et al., 1999; Schreiber, 2003; Gascoigne and Lipcius, 2004). Positive density dependence might also arise in populations of ecosystem engineers, organisms that significantly modify the surrounding abiotic and biotic environment, as the population size must be sufficiently large to generate required environmental change for population persistence (Byers et al., 2006; Cuddington et al., 2009). If the Allee effect is strong enough, at low population sizes the population will experience negative growth rates, and ultimately extinction (Courchamp et al., 1999). This leads to a critical population size required for population persistence; below this critical threshold, the population will decline to extinction, while above this threshold the population will persist. Knowledge of this critical threshold is thus of particular importance in exploited or degraded populations where there is an interest in population restoration or conservation (Courchamp et al., 2008).

One ecosystem engineer of particular restoration importance is the Eastern oyster, *Crassostrea virginica*. This species inhabits thousands of miles of coastline, and provides valuable ecosystem services, including commercial harvest, water filtration, seashore stabilization, erosion protection, and habitat and predator refuge for a variety of organisms (Coen et al., 2007; Grabowski et al., 2012). In these oyster populations, individuals aggregate into large, complex reef structures. Reefs are composed of living oysters, as well as oyster shell that remains following natural mortality. Shell provides solid substrate on which new oyster larvae can attach, increasing larval survival by providing shelter from predators and preventing burial in sediment (Rothschild et al., 1994; Mann and Powell, 2007). Reefs also increase growth and survival of adult oysters by increasing water filtration, buffering against hypoxic events, and increasing food availability through increased current speeds (Lenihan et al., 1996; Lenihan and Peterson, 1998; Lenihan, 1999; Bartol et al., 1999; Schulte et al., 2009).

Globally, oysters have experienced severe population declines due to decades of overfishing, coastal development, and pollution (Airoldi and Beck, 2007; Beck et al., 2011). Particularly damaging has been the use of destructive fishing practices that not only remove older, more fecund individuals, but also destroys the reef structure and available substrate that is necessary for recruitment and persistent populations (Rothschild et al., 1994). Additionally, the emergence and increasing prevalence of two protozoan diseases, MSX and Dermo, along the eastern United States in the mid to late 1900s has contributed to population declines (Hofmann et al., 2009; Carnegie and Burreson, 2011). Along the eastern coast of the United States, many native *C. virginica* populations have been reduced to less than 15% of their historic population sizes, with an associated decline in substrate availability and integrity (Rothschild et al., 1994; Beck et al., 2011; zu Ermgassen et al., 2012).

Demographic modeling has shown that positive feedbacks between living oysters and shell substrate can lead to thresholds between population persistence and extinction, as well as possible alternative stable states (Jordan-Cooley et al., 2011; Nystrom et al., 2012; Housego and Rosman, 2016). In systems with alternative stable states, restoration becomes particularly challenging as transitions between desired and undesired states can occur through sudden, often unpredictable, regime shifts, and successful restoration often requires the conditions of the system be returned to levels more extreme than those of the original system (Beisner et al., 2003; Scheffer et al., 2001; Scheffer and Carpenter, 2003; Hastings and Wysham, 2010). In oysters, the desirable state consists of a healthy, abundant population of oysters on high-relief reefs, while the undesirable state is heavily degraded, with low or zero population sizes.

Empirical data and restoration efforts also supports these ideas (Powell et al., 2009a,b; Schulte et al., 2009). For example, Powell et al. (2009b) analyzed a time series of *C. virginica* populations in Delaware Bay from 1953-2006, and found that the population persisted for extended periods of time in two distinct states, one of high abundance, and one of low abundance. Additionally, Schulte et al. (2009) showed that the success of restoration of *C. virigincia* populations in the Chesapeake Bay was influenced significantly by the vertical height of the reef. Locations restored with high vertical reefs had greater oyster densities and were likely to persist, while populations restored with low vertical reefs had low oyster densities and were predicted to decline to extinction within a handful of years. This result suggests a critical threshold of reef height required for persistence.

Given the possible existence of a threshold between population persistence and extinction, it is important to understand the shape of this threshold in size-structured populations, and what restoration actions can be taken to push a population from an extinction trajectory to a persistent trajectory. Restoration efforts in oyster populations generally consist of (i) stock enhancement, i.e. supplementing existing populations with additional oyster spat reared in the lab, or oysters grown in aquaculture-like “oyster gardens”; (ii) habitat enhancement, i.e. adding recycled shells or artificial reef structures to increase the availability of substrate; or (iii) a combination of stock and habitat enhancement (Brumbaugh and Coen, 2009). While there has been successful restoration of some *C. virginica* populations along the mid-Atlantic US coast (Taylor and Bushek, 2008; Powers et al., 2009; Schulte et al., 2009; Puckett and Eggleston, 2012; Lipcius et al., 2015), there are concerns about the efficacy of restoration actions, particularly given high disease prevalence, and there is no current agreement on the best means for achieving success (Mann and Powell, 2007; Kennedy et al., 2011; Geraldi et al., 2013; but see Baggett et al., 2014 and Lipcius et al., 2015).

Here, we use *C. virginica* as a model species to investigate the impact of positive density dependence on population dynamics and restoration actions. Specifically, we are interested in understanding the properties of the threshold between population persistence and extinction. We extend a closed, size- and age-structured integral projection model (IPM) developed in Moore et al. (2016) to include a positive feedback between the establishment of new oyster larvae and shell substrate. We use this model to address several questions. First, we explore properties of the threshold and introduce a general analytic method for approximating the infinite dimensional threshold surface. We next investigate the affect of population size structure on the threshold surface, and the ultimate fate of the population. Finally, we assess the relative effectiveness of two restoration actions, namely stock enhancement using oysters of different ages, or habitat enhancement, at restoring a population declining toward extinction such that the population persists.

## METHODS

### Model

We extend an age- and size-structured integral-projection model (IPM) developed in Moore et al. (2016). Fig. 1 shows a simplified representation of the full model. Briefly, let *n_a_*(*x*, *t*)*dx* be the density of age *a*, size *x* oysters at time *t*, with *x* measured as the shell length of an oyster in mm. Oysters of size *x* will survive to the next time step and grow to size *y* according to age- and size-specific survival and growth kernels, *S_a_*(*x*) and *G_a_*(*y*, *x*), respectively. The fecundity kernel, *F_a_*(*y*, *x*, *H*(*t*)), represents the density of size *y* recruits produced by an adult of age *a* and size *x*. The fecundity kernel also depends upon, *H*(*t*), the m^2^ of substrate available at time *t*. Though adult oysters are also positively affected by the amount of substrate, this effect is due largely to the physical location of the oyster within the reef structure (Lenihan and Peterson, 1998; Lenihan, 1999), and is thus more indirect than the effect on the oyster larvae. Thus, we chose to focus solely on the positive effect of substrate levels on recruitment. The dynamics of the population are expressed as

**Figure 1.**
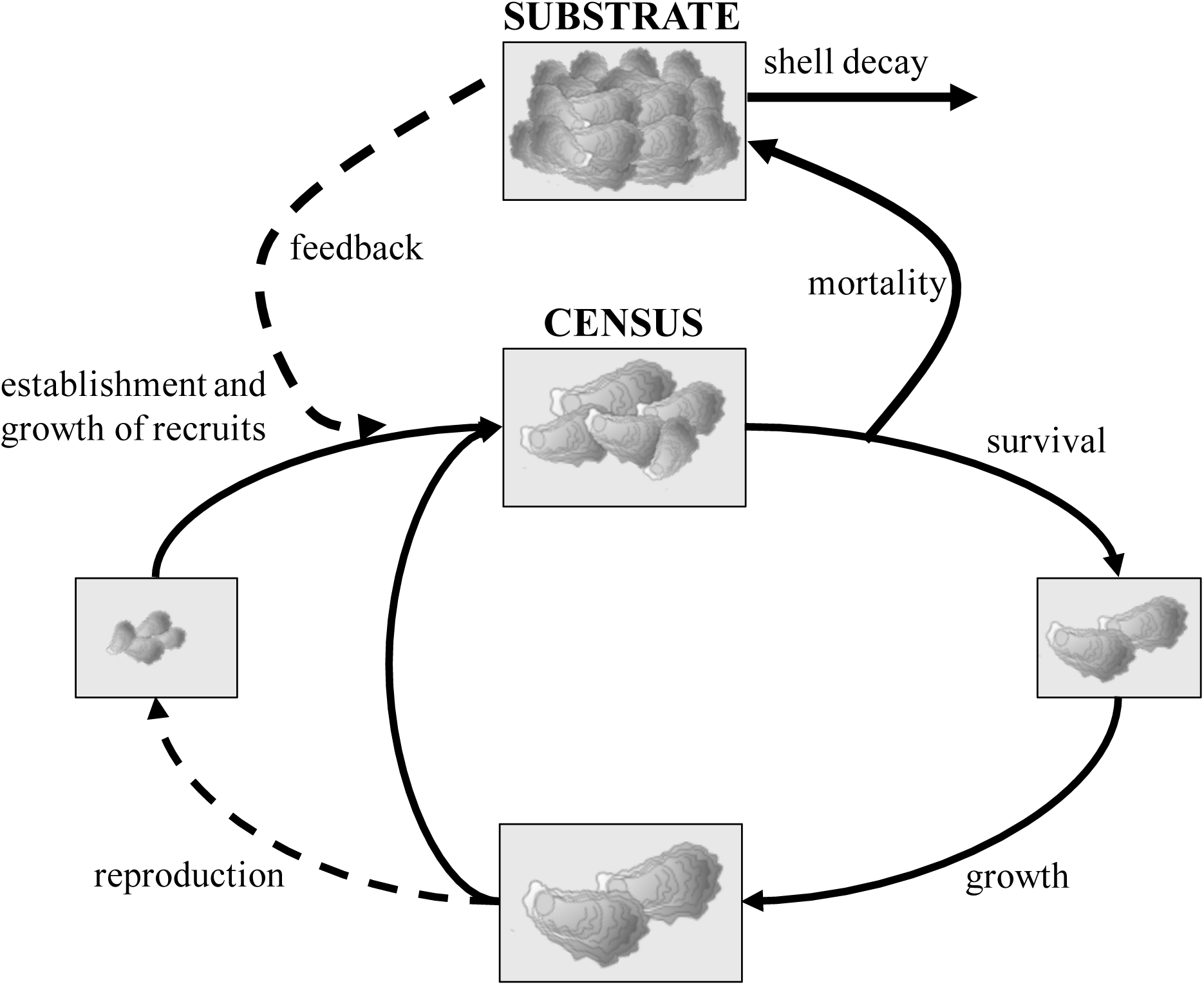
Representation of model. Census occurs immediately following summer recruitment. Oysters then experience mortality, with dying oysters converted to a surface area of substrate. Surviving oysters grow before reproducing. Following reproduction, new oyster recruits experience a separate growth event before joining existing oysters immediately prior to the next census. The number of recruits that successfully establish depends upon the degree of local retention, and the feedback with substrate.

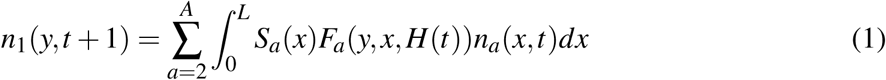

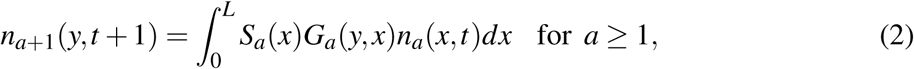

where *A* is the maximum age of an individual, and *L* is the maximum size of an individual. To avoid artificial eviction of individuals growing larger than the maximum size, we include a discrete size class for individuals of size *x* > *L*, with kernels for survival and fecundity set equal to kernels for individuals of size *x* = *L* (Moore et al., 2016; Williams et al., 2012). This discrete class is given by

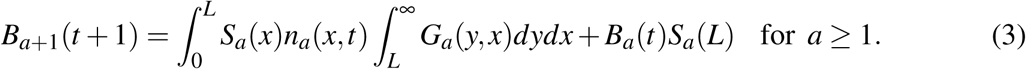

with *B*_1_ = 0. We assume that available substrate is equal to the total surface area of shell in the population, with substrate dynamics given by

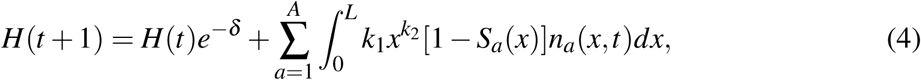

where *δ* is the decay rate of shell (per year), and *k*_1_ and *k*_2_ are scaling parameters that convert the number of size *x* individuals that experienced mortality to m^2^ of surface area. Though oyster larvae are not restricted to settlement on dead conspecifics (Nestlerode et al., 2007), we choose the surface area of shell to better reflect units most relevant to restoration practices.

In Pamlico Sound, spawning of *C. virginica* is protracted, with a primary spawning and settlement peak in May and June, and a secondary spawning and settlement peak in July and August (Ortega and Sutherland, 1992; Mroch et al., 2012). For simplicity, we model reproduction as occurring once at the end of the spring, with census occurring immediately thereafter. Between census and reproduction, adult oysters experience mortality, then grow from their current size *x* to their final end-of-year size, *x′*. Reproduction then occurs according to a size-specific fecundity relationship, *f* (*x′*, *H*(*t*)). This fecundity relationship is composed of two parts: (i) the number of new oyster larvae that are produced, survive, and settle in the natal population; and (ii) the feedback between available substrate and the fraction of larvae that are able to successfully establish. We assume the number of eggs produced is dependent upon the size of the parent, and that all oyster recruits are created by adults of the local population (i.e. there is no immigration). The fecundity relationship is given by

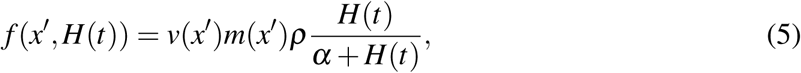

where *v*(*x′*) is the proportion of size *x′* individuals in the population that are female, *m*(*x′*) is the number of eggs produced by a size *x′* individual, *ρ* is the maximal local retention of oyster larvae (e.g. the proportion of eggs that survive and settle in the natal population), and *H*(*t*)/(*α* + *H*(*t*)) represents the positive feedback between available substrate and recruitment. The sizes of the newly recruited oysters are assumed to be normally distributed with density *z*(*y*). Thus, the overall fecundity kernel is expressed as

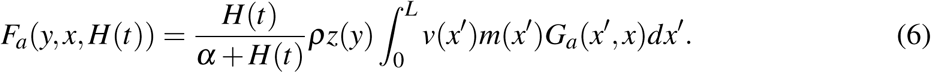

While there is evidence that negative density dependence is also important in oyster systems, potentially leading to alternative stable states (Knights and Walters, 2010; Puckett and Eggleston, 2012; Jordan-Cooley et al., 2011; Powell et al., 2009b), here we are only interested in what determines population persistence versus extinction, rather than the properties of the system at some stable carrying capacity. As such, we restrict our focus to investigating the effects of positive density dependence, rather than the effects of both positive and negative density dependence.

### Data

We estimated site-specific kernels for growth, survival, and fecundity using data collected from the West Bay *C. virginica* population in Pamlico Sound, North Carolina. A full description of the methods is provided in Mroch et al. (2012) and Puckett and Eggleston (2012, 2016). Briefly, data to estimate growth and survival kernels were obtained by deploying 15 replicate settlement trays at a restored oyster reef protected from harvest. On each tray, individual settlers were marked and growth and mortality tracked from June 2006 to October 2008. To estimate size-specific per capita fecundity, oysters were collected from the reef and brought back to the lab to determine the total egg content of each oyster following the general procedures of Cox and Mann (1992).

### Statistical fitting

*Growth and survival kernels*.— To estimate the growth kernel, *G_a_*(*y*, *x*), we fit a linear regression of the log change in size from time *t* to *t* + 1 against the size at time *t*, assuming constant variance across all ages and sizes. Fitting growth in this way ensures non-negative changes in size, which is important for describing oyster growth (Moore et al., 2016). We fit the survival kernel, *S_a_*(*x*), using logistic regression of survival between years. We assume that mortality is age- and size-dependent, with larger, older oysters more susceptible to diseases, and juveniles more susceptible to predation. While we do not measure these effects explicitly, we assume these processes are captured implicitly in the field data.

*Fecundity kernel*.— We estimate the size-specific number of eggs produced, *m*(*x′*), with a scaling relationship. Using the estimated number of eggs produced during May 2007 and May 2008 (Mroch et al., 2012), we fit a linear relationship between the log number of eggs and log oyster size. Since oysters are protandric hermaphrodites, beginning life as male and switching to female at larger sizes (Galtsoff, 1964), we expect a higher proportion of females at larger sizes. We thus estimate the size-specific proportion of females in the population, *v*(*x′*), using a linear regression of the proportion of females in the population against size, using data from May 2007 and May 2008 (Mroch et al., 2012). After fitting the model, we bound the function such that any negative value was set equal to zero, while any value greater than one was set equal to one. However, model results are not highly sensitive to the form chosen for *v*(*x′*). We estimate the size distribution of new recruits, *z*(*y*), with a normal distribution, using the mean and standard deviation of measured recruit sizes from August 2006 and August 2007 (Puckett and Eggleston, 2012).

Local retention, *ρ*, depends upon factors such as fertilization success, survival during the pelagic larval stage, predation, and local dispersal and transport processes. Here, we parameterize *ρ* using results of a coupled hydrodynamic and particle tracking simulation presented in Puckett et al. (2014). Briefly, larval dispersal was simulated over a 21 day period, whereby a daily mortality rate of 0.2 per day was applied. After 14 days, larvae were assumed to settle if located within the reef polygon. Local retention was estimated as the proportion of larvae spawned from a reef that settled within their natal reef. We also consider the case when local retention is low. For this case, we set *ρ* to 50% above the minimum value of local retention that still yields a positive equilibrium (see *Methods: Model analysis*).

Finally, to estimate the *α* parameter of the positive feedback function, we solve for *α* using Eqn. 5 multiplied by *n*(*x′*). Thus,

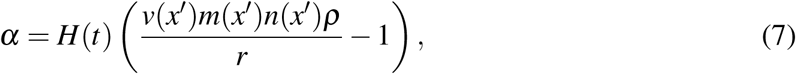

where *r* is the observed, size-independent number of recruits. We obtain estimates of *r*, *n*(*x′*), and *H*(*t*) from Puckett and Eggleston (2012). Estimates of *v*(*x′*), *m*(*x′*), and *ρ* are as described above.

*Substrate dynamics*.— We obtain estimates of the substrate decay rate, *δ*, from Wilberg et al. (2013). For *k*_1_ and *k*_2_, the scaling parameters between oyster length and surface area, we used the scaling relationship given in Galtsoff (1966).

### Model analysis

To understand the dynamics of the model, we first assess model behavior by analytically determining the location of the threshold surface dividing regions of population persistence from regions of population extinction. We then conduct an elasticity analysis to determine how the location of this threshold surface is affected by changes in model parameters. Finally, we assess several restoration scenarios using analytic approximations and numerical simulations.

*Model behavior*.— The theory developed in Schreiber (2004) applies to the discretization of the IPMs which are used for all our numerical work. This theory characterizes the dynamical behavior of the model using the dominant eigenvalue of the model at low densities, *λ*_0_, and the dominant eigenvalue at high densities, *λ*_∞_. Using these eigenvalues, there are three possible dynamics: (i) asymptotic extinction for all initial densities when *λ*_∞_ < 1; (ii) unbounded growth (persistence) for all non-zero initial densities when *λ*_0_ > 1; and (iii) the existence of a co-dimension one threshold surface such that initial conditions below this surface lead to extinction, while initial conditions above this surface lead to unbounded growth (persistence). For the parameters considered here, the model always exhibits the third behavior. Moreover, as we will show, there is a unique unstable equilibrium on this threshold surface. We will use linearizations at this unstable equilibrium to gain insights into the geometry of the threshold surface.

At the unstable equilibrium on the threshold surface, the dominant eigenvalue, *λ*, of the demographic transition operator equals one. Let *ρ̂* be the value of *ρ* such that *λ* = 1 in the linear model (i.e. when *f*(*x′*) = *v*(*x′*)*m*(*x′*)*ρ̂*), and *n̂_a_*(*x*) be the stable size- and age-distribution when *λ* = 1 in the linear model. To solve for the equilibrium in the non-linear model with positive feedbacks, we equate the fecundity term in the non-linear model with the fecundity term in the linear model at equilibrium: 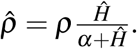 Solving for the equilibrium level of substrate, *Ĥ*, yields

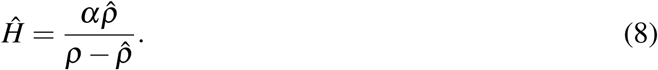

Note that for a positive equilibrium, *ρ* > *ρ̂*.

We then solve for the total equilibrium number of oysters using Eqn. 4 and evaluating *n_a_*(*x*, *t*) at *n̂_a_*(*x*)*N̂*, where *N̂* is the total number of oysters at equilibrium across all sizes and ages. Thus,

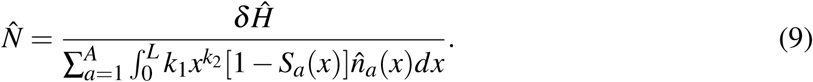

To investigate model behavior around the unstable equilibrium, we run simulations of population trajectories using two initial size and age distributions: (i) the equilibrium size and age distribution, *n̂_a_*(*x*); and (ii) a harvested age and size distribution, set by truncating and re-normalizing the equilibrium size and age distribution such that all oysters ≥ 75 mm were removed from the population. This latter distribution roughly approximates a population experiencing severe harvesting pressure. For each of the two initial size and age distributions, we numerically estimate the separatrix of total oyster numbers and total substrate above which the population would persist, and below which the population would decline to extinction. We do this using a bisection search algorithm. Briefly, we first set the initial amount of substrate, *H*_0_. We then determine an initial value of total oysters, *c*_1_, such that a population beginning at this value is above the threshold surface and persists, and an initial value of total oysters, *c*_2_, such that a population beginning at this value is below the threshold surface and goes extinct. We then run the simulation with an initial total oyster number equal to *c*_3_ = (*c*_1_ + *c*_2_)/2. If this new population is above the threshold surface, in the next simulation we set the new total oyster number equal to *c*_4_ = (*c*_2_ + *c*_3_)/2, otherwise we set the new total oyster number equal to *c*_4_ = (*c*_1_ + *c*_3_)/2. We repeat this process *k* steps until *c_k_* is above the threshold surface and *c_k_* – *c*_*k*–1_ < 0.001*N̂*. We repeat this process across a range of values of *H*_0_ near *Ĥ* and for each of the two initial size and age distributions.

*Elasticity analysis*.— Given that many parameters of the model are highly uncertain, and potentially variable across space and time, we compute the elasticity of the equilibrium to local retention, *ρ*, the shape parameter of the feedback function, *α*, and the substrate decay rate, *δ*. These elasticities indicate the percentage change in the equilibrium values given a 1% change in the parameter. Let **x** = (**n**, *H*)^T^, **n** = (*n*_1_(*y*), *B*_1_, ⋯, *n_A_*(*y*), *B_A_*), *A*(*H*, *θ*) be the full demographic operator (Eqns.1–3), *h*(**x**, *θ*) be the dynamics of the substrate (Eqn.4), and *θ* be the parameter of interest. Then, the sensitivity of the equilibrium population densities and substrate level to *θ* is

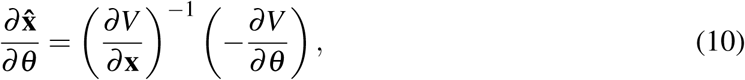

where

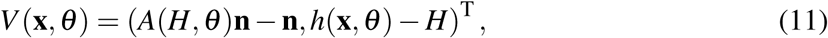

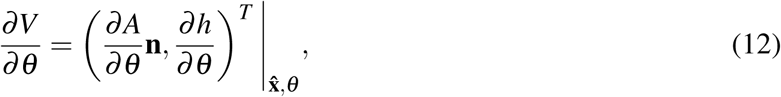

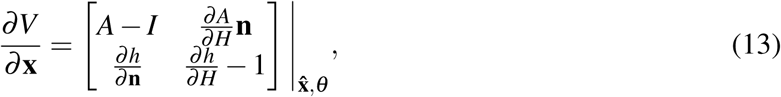

and *I* denotes the identity operator. The elasticity of the population sizes and substrate level to *θ* is then given by

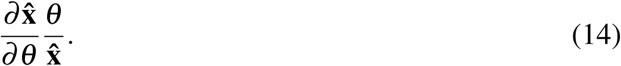

Appendix S1 gives the specific equations used to calculate the elasticity of the equilibrium population densities to *δ*, *ρ*, and *α*.

*Restoration scenarios and analytic approximations*.— Restoration of an oyster population is desirable if the population lies below the threshold surface and is heading toward extinction. Successful restoration is then defined here as restoration actions that push the population across the threshold surface such that the population will theoretically increase toward infinity. Here, we consider two types of one-shot restoration actions: (i) the addition of cohorts of oysters of a single age, *a*; and (ii) the addition of shell substrate. Since restoration actions require significant amounts of time and money, we are interested in determining the minimum amount of either oysters or substrate that will push the population across the equilibrium threshold surface. We approach this question using an analytic approximation for oyster additions, and numerically for both oyster or substrate additions.

First, we use Eqn.13 to investigate the linearization of the threshold surface around the unstable equilibrium. The left dominant eigenvector of this operator gives the direction perpendicular to the threshold surface, and thus indicates the relative amount of oysters of a particular age and size that should be added to minimize the distance between the threshold surface and a point below the threshold surface (Appendix S2). Since we are interested in adding cohorts of oysters of a particular age to better reflect restoration practices, we use the relationship between the left eigenvector, the unstable equilibrium, and the size distribution of age *a* oysters to analytically approximate the total number of age *a* oysters required to cross the threshold surface (Appendix S3). We determined the size distribution of age *a* oysters by applying the growth and survival kernels to the initial distribution of age 1 recruits, *z*(*y*), and re-normalizing the distribution after each time step.

To determine whether the analytic approximations of the required oyster numbers work well, we also simulate restoration actions numerically to determine the minimum number of age *a* oysters, or the minimum amount of substrate required to push the population over the threshold surface, assuming a one-time addition of either substrate or oysters of age *a*. To numerically determine the minimum number of oysters of age *a* or substrate required, we use the bisection search algorithm described in *Methods: Model analysis*. For simulations of oyster additions, we assume the population starts with no living oysters, and no available substrate. This assumes a worst-case scenario in which an oyster population once existed, but became degraded to the extent that no oysters or shell remain. For substrate additions, we assume the population starts with no available substrate, and a number of oysters equal to 10% above or 10% below *N̂*. We modeled the size distribution of existing oysters using either the equilibrium size distribution, or the harvested size distribution.

In all model analyses, we discretized the integral operators using the midpoint rule with 250 equally sized bins from size 0 to 250 mm, for each age class from 0 to 10 years. We ran all simulations for 150 time steps. Model implementation and data analysis were conducted with R (R Core Team, 2015).

## RESULTS

### Statistical fits

A total of 590 oysters were observed for approximately 2 years post-settlement from June 2006 to October 2008. Measured oyster sizes ranged from 5.4 mm to 86.1 mm, while oyster ages ranged from 62 days to 2.3 years. Oyster sizes observed from quadrat sampling (to obtain individuals to estimate per-capita fecundity) ranged from 6-124 mm on substrate that was 3-5 years old over the course of the study (Mroch et al., 2012; Puckett and Eggleston, 2012). In the model, we extrapolated both size and age to span a biologically realistic range of values, allowing size to vary from 0 mm to 250 mm, and age to vary from 0 days to 10 years.

While previous work has shown the importance of including both age- and size-structure in models of oyster populations (Moore et al., 2016), the limited time-frame of our data led to poor fits for growth and survival functions when including both age and size. We thus fit growth and survival functions using only size, but also included a maximum age of survival, *A* = 10, beyond which no oysters survive, regardless of size. The final growth function shows a negative relationship between the log change in size and size of an oyster. When translated to the relationship between size at time *t* + 1 and size at time *t*, this led to a growth function in which oyster growth slowed as size increased (Fig. 2A). Oyster survivorship increased as a function of size (Fig. 2B).

**Figure 2.**
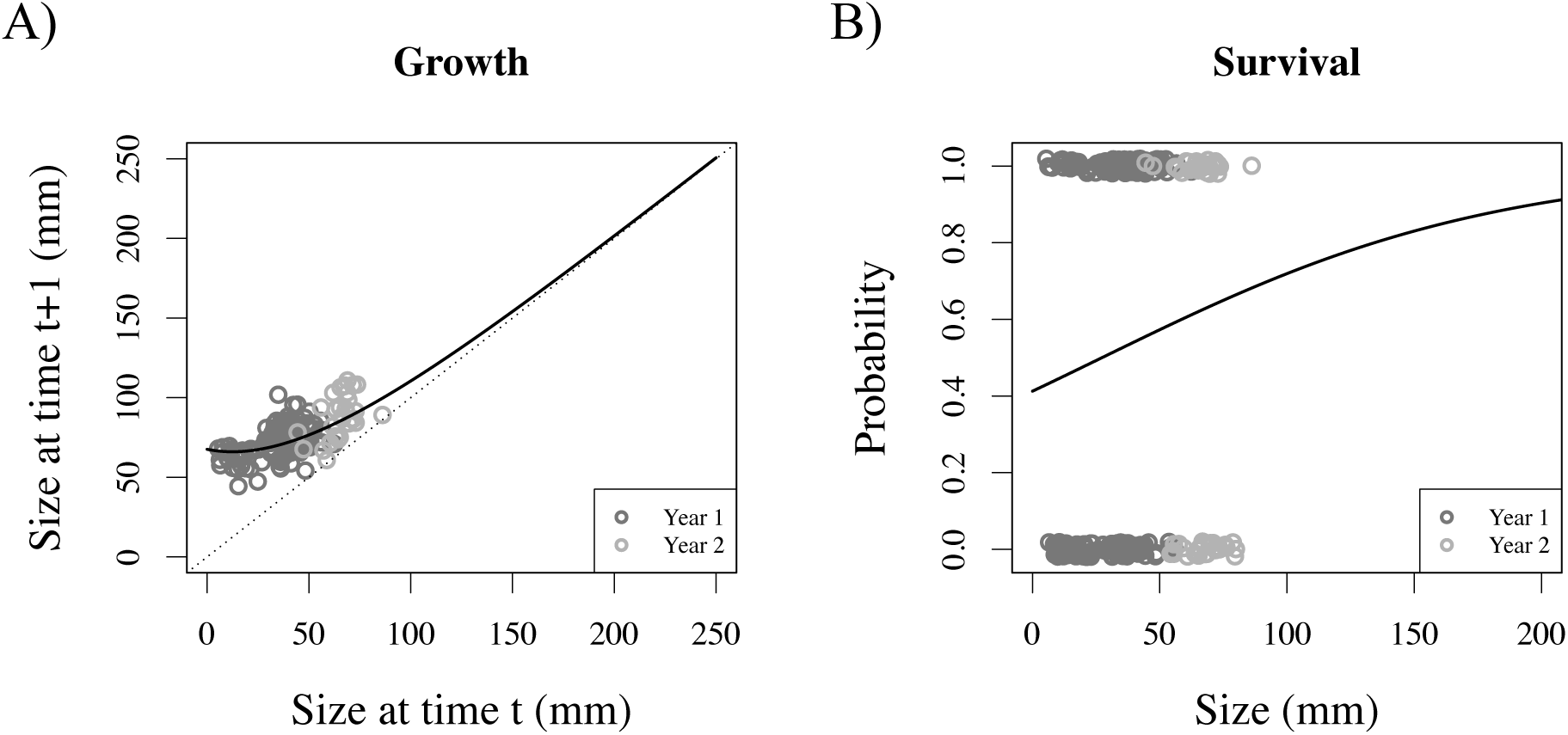
Growth and survival functions. Statistical fits of size-dependent growth (A) and survival (B) functions. A) Growth functions are fit using linear regression on the log change in size against size, then translated to generate the relationship between size at time *t* + 1 and size at time *t*. The dotted black diagonal line is the 1:1 line. B) Survival functions are fit using linear regression of survival between time points. All functions are extrapolated past the collected data (gray points) to the minimum (0 mm) and maximum (250 mm) sizes. Model parameters are given in Table 1.

**Table 1:**
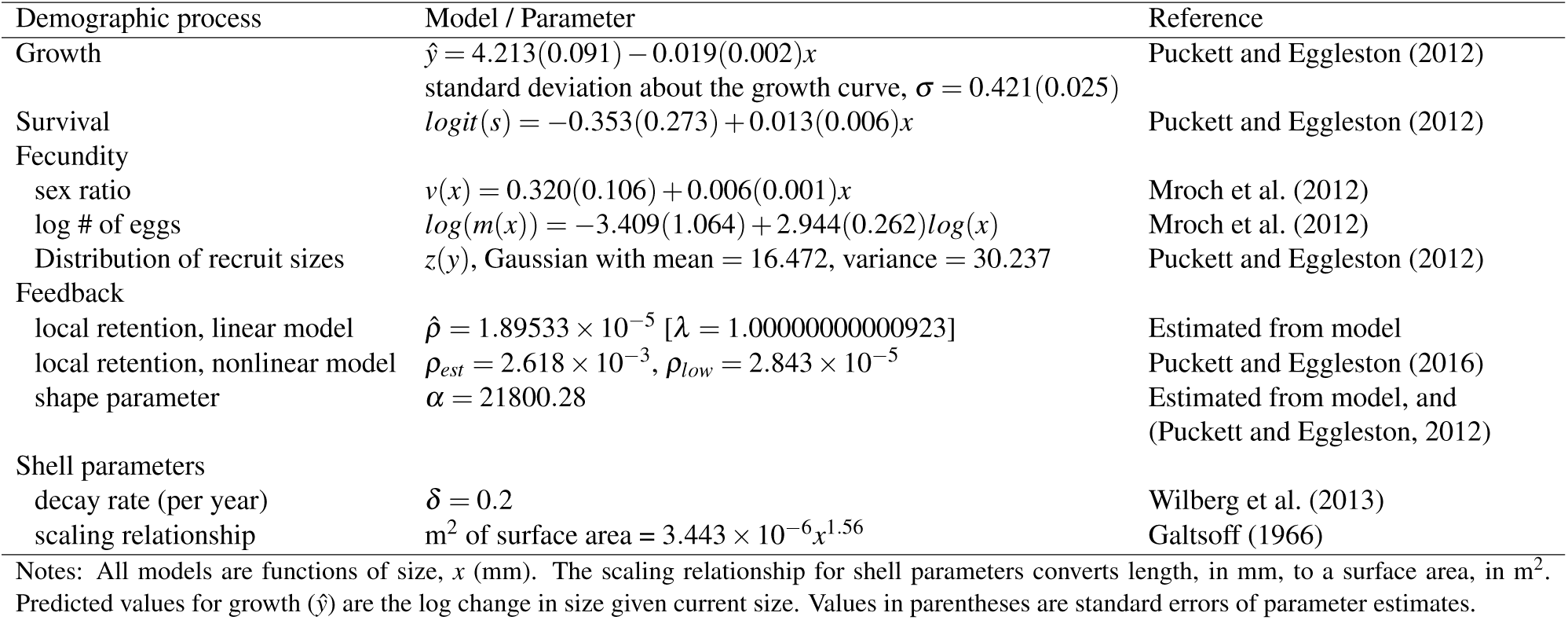
Demographic functions and parameter estimates. Statistical models and parameter estimates for the size- and age-structured model used to describe *C. virginica* demography.

The proportion of females in the population increased as a function of size (Fig. 3A), while the log number of eggs increased linearly as a function of log female size (Fig. 3B). The size distribution of new recruits was normally distributed with mean = 16.47 mm, and sd = 5.50 mm (Fig. 3C).

**Figure 3.**
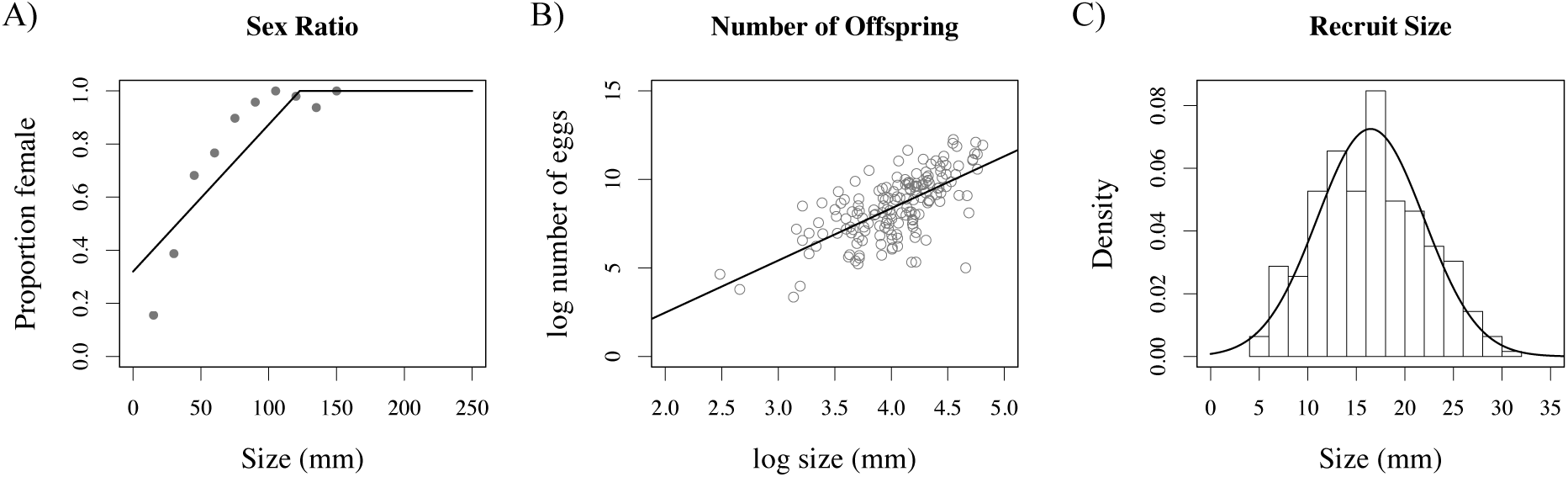
Fecundity functions. A) The proportion of females as a function of size. B) The log number of eggs produced as a function of log female size. C) The distribution of offspring size, fit to August 2007 and 2008 recruit sizes. Data for (A) and (B) from Mroch et al. (2012), and data for (C) from Puckett and Eggleston (2012). Parameters of all model fits are given in Table 1.

We set the degree of local retention *ρ*_est_ = 2.617898 × 10^−3^, using estimates from Puckett and Eggleston (2016). We also investigated model results when the degree of local retention was low (for instance, if environmental conditions changed such that recruit survivorship decreased), and set *ρ*_low_ = 2.842991 × 10^−5^. This value was chosen to be equal to 50% above *ρ̂*, the minimum value of local retention that still yields a positive equilibrium. To find the unstable equilibrium, we used a value of *ρ̂* = 1.89532724809747 × 10^−5^, which yielded a long-term population growth rate of *λ* = 1.00000000000923 in the linear model.

To obtain an estimate of *α*, we used *H* = 4134.5 m^2^ (Puckett and Eggleston, 2012), and a total population size of *N* = 3, 583, 233.333 oysters (Puckett and Eggleston, 2016). We multiplied the total population size by size-frequency data of oysters at this location to obtain an estimate of the size distribution *n*(*x*) (Puckett and Eggleston, 2016). To estimate *r*, we used the number of observed recruits in August 2006 to obtain an estimate of *r* = 4, 266, 804 new recruits (Puckett and Eggleston, 2012). Since *r*, as estimated in Puckett and Eggleston (2012), measures the overall number of recruits, it is possible that this value includes immigrant recruits produced from outside the local population. However, as the West Bay site is relatively isolated from other nearby oyster reefs, the input of oyster recruits from external sources is likely small (Puckett and Eggleston, 2016). Plugging these values, as well as *ρ*_est_ and size-specific sex ratios and eggs as described above into Eqn. 7 yielded a value of *α* = 21, 800.28.

Wilberg et al. (2013) gives a range of shell decay rates from 0.05-0.4 per year. We used a mid value from this range and set *δ* = 0.2. We also explored dynamics for values of *δ* at the extremes of this range. Galtsoff (1966) give the scaling relationship between oyster length, in cm, and surface area, in cm^2^ as 1.25(L_cm_)^1.56^. Converting to mm and m^2^, respectively, yields *k*_1_ = 3.443 × 10^−6^ and *k*_2_ = 1.56, and thus m^2^ = 3.443 × 10^−6^(L_mm_)^1.56^.

All demographic functions and parameter estimates are given in Table 1.

### Model analysis

*Model behavior*.— Using the parameter values in Table 1, the unstable positive equilibrium is (*N̂*, *Ĥ*) = (20, 928.69 oysters, 158.98 m^2^) when *ρ* is high (*ρ*_est_), and (*N̂*, *Ĥ*) = (5, 739, 643.58 oysters, 43, 601.02 m^2^) when *ρ* is low (*ρ*_low_). For populations beginning at the equilibrium size distribution (Fig. 4B), the separatrix passes through the unstable equilibrium (Fig. 4A), while for populations beginning at the harvested size distribution (Fig. 4C), the separatrix is greater than that of the equilibrium size (Fig. 4A). Above the harvested separatrix (Fig. 4A, region I), populations beginning at either initial size distributions will increase to infinity, while below the equilibrium separatrix (Fig. 4A, region III) populations beginning at either the equilibrium or the harvested size distribution will decline to extinction. However, there exists a large region between the two separatrices (Fig. 4A, region II), where populations beginning at the equilibrium size distribution will increase to infinity, but populations beginning at the harvested size distribution will decline to extinction, even if the population begins above (*N̂*, *Ĥ*). Additionally, populations beginning at the harvested size distribution will often exhibit complex, oscillatory behavior during the first 10-12 years of the simulation, regardless of whether they are declining toward extinction or increasing toward infinity.

**Figure 4.**
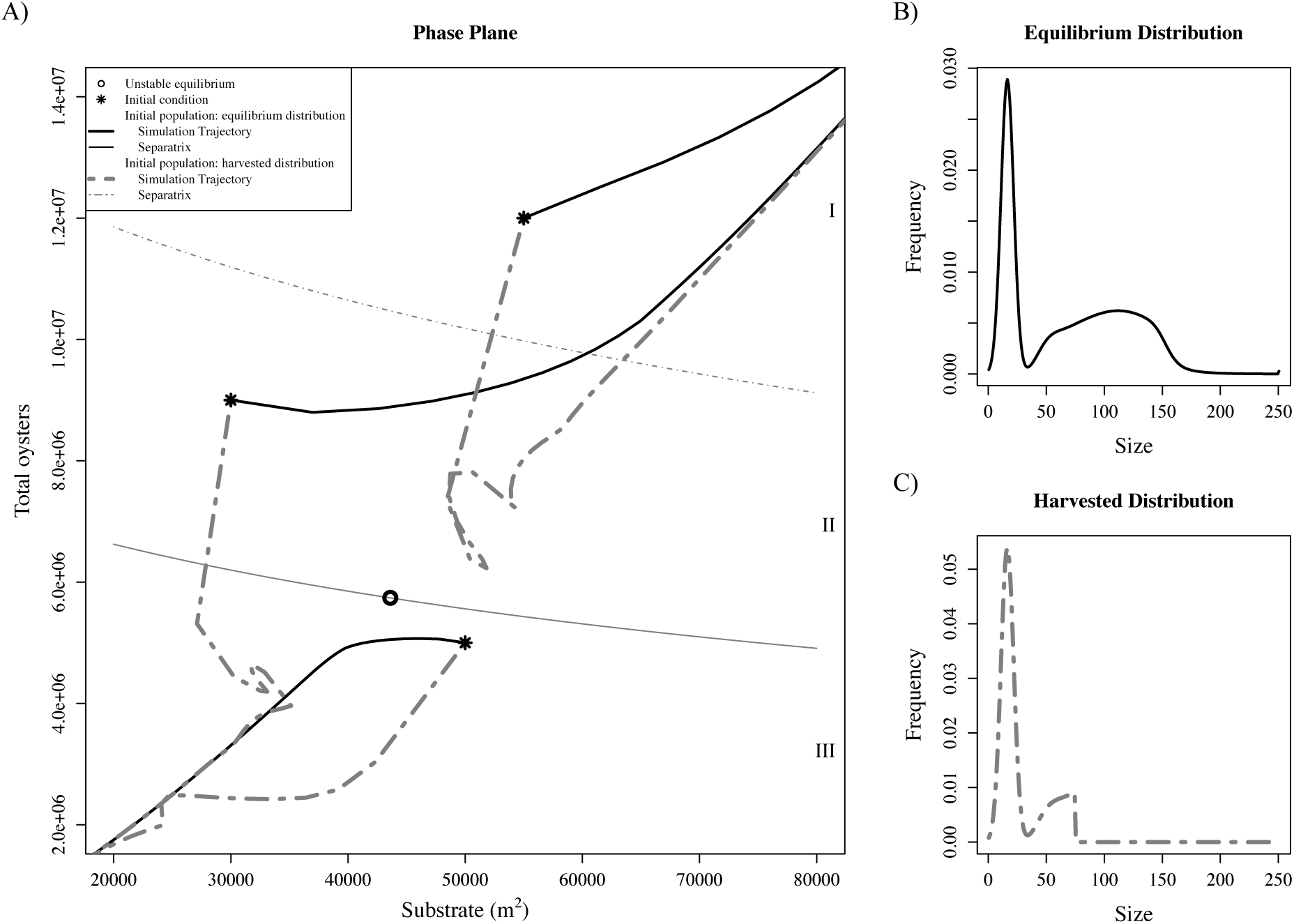
Population trajectories and initial size distributions. A) Population trajectories in the absence of restoration for *ρ*_low_, with remaining parameters given in Table 1. Populations begin at either the equilibrium size distribution (B; thick black lines), or a harvested size distribution (C; thick gray lines). Black stars indicate initial conditions, while the black circle indicates the unstable equilibrium, (*N̂*, *Ĥ*). Thin lines give the projected separatrix for populations beginning at the equilibrium size distribution (solid black), or the harvested size distribution (dotted gray). Above the harvested separatrix (region I), populations beginning at either size distribution will increase to infinity, while below the equilibrium separatrix (region III), populations beginning at either size distribution will decline to extinction. Above the equilibrium separatrix but below the harvested separatrix (region II), populations beginning at the equilibrium size distribution will increase to infinity, while populations beginning at the harvested size distribution will decline to extinction. B) The size distribution of the population at equilibrium (when *λ* = 1 in the linear model). C) The size distribution of a harvested population, obtained by truncating the equilibrium size distribution such that all oysters ≥ 75 mm are removed from the population.

*Elasticity analysis*.— Increasing the feedback parameter, *α*, will increase **x̂**, while increasing local retention, *ρ*, will decrease **x̂** (Fig. 5). Increasing the substrate decay rate, *δ*, will increase *N̂* and decrease *Ĥ*, though the effect on *Ĥ* is much smaller than the effect on *N̂* (Fig. 5). Of the three parameters, changes in local retention *ρ* has the largest impact on the threshold surface, while changes in *δ* has the smallest impact on the threshold surface. The effect on **x̂** to changes to *ρ* is reduced when *ρ* is high, while the effect on **x̂** to changes to *δ* is reduced when *δ* is low (not shown).

**Figure 5.**
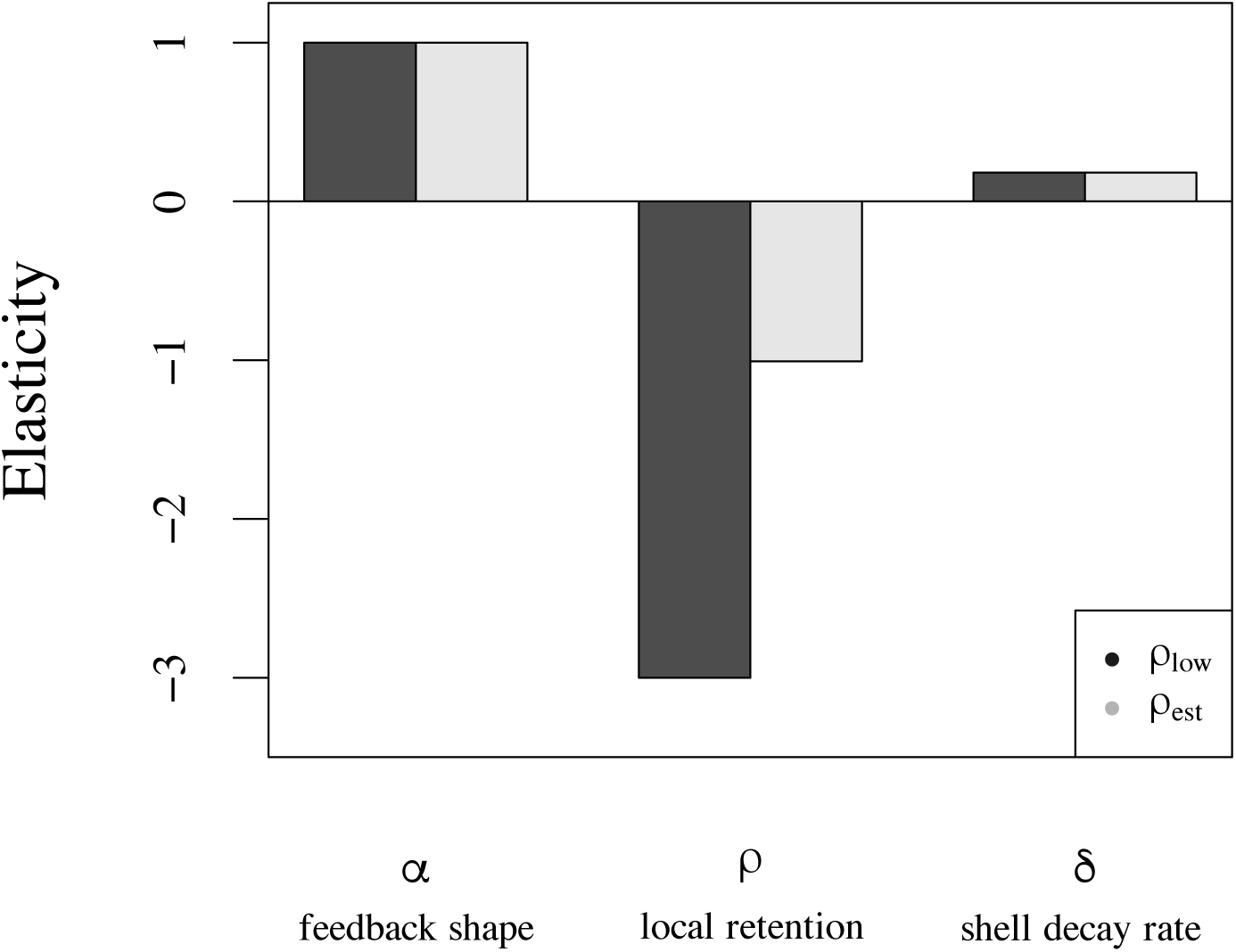
Elasticity analysis. Elasticity of equilibrium oyster numbers for *ρ*_low_ (dark bars) and *ρ*_est_ (light bars), with remaining parameters given in Table 1. The elasticity of equilibrium substrate levels is the same as the elasticity of equilibrium oyster numbers for *α* and *ρ*. The elasticity of *Ĥ* to *δ* is equal to −2.4 × 10^−11^ for *ρ*_low_ and −8 × 10^−12^ for *ρ*_est_.

*Restoration scenarios*.— Fig. 6 shows the normalized size distributions of age *a* oysters. Using these distributions and the methods presented in Appendix S3, we analytically approximate the total number of oysters of a particular age cohort that are required to cross the threshold surface. These results, as well as the results of the numerical simulations, are shown in Fig. 7A.

**Figure 6.**
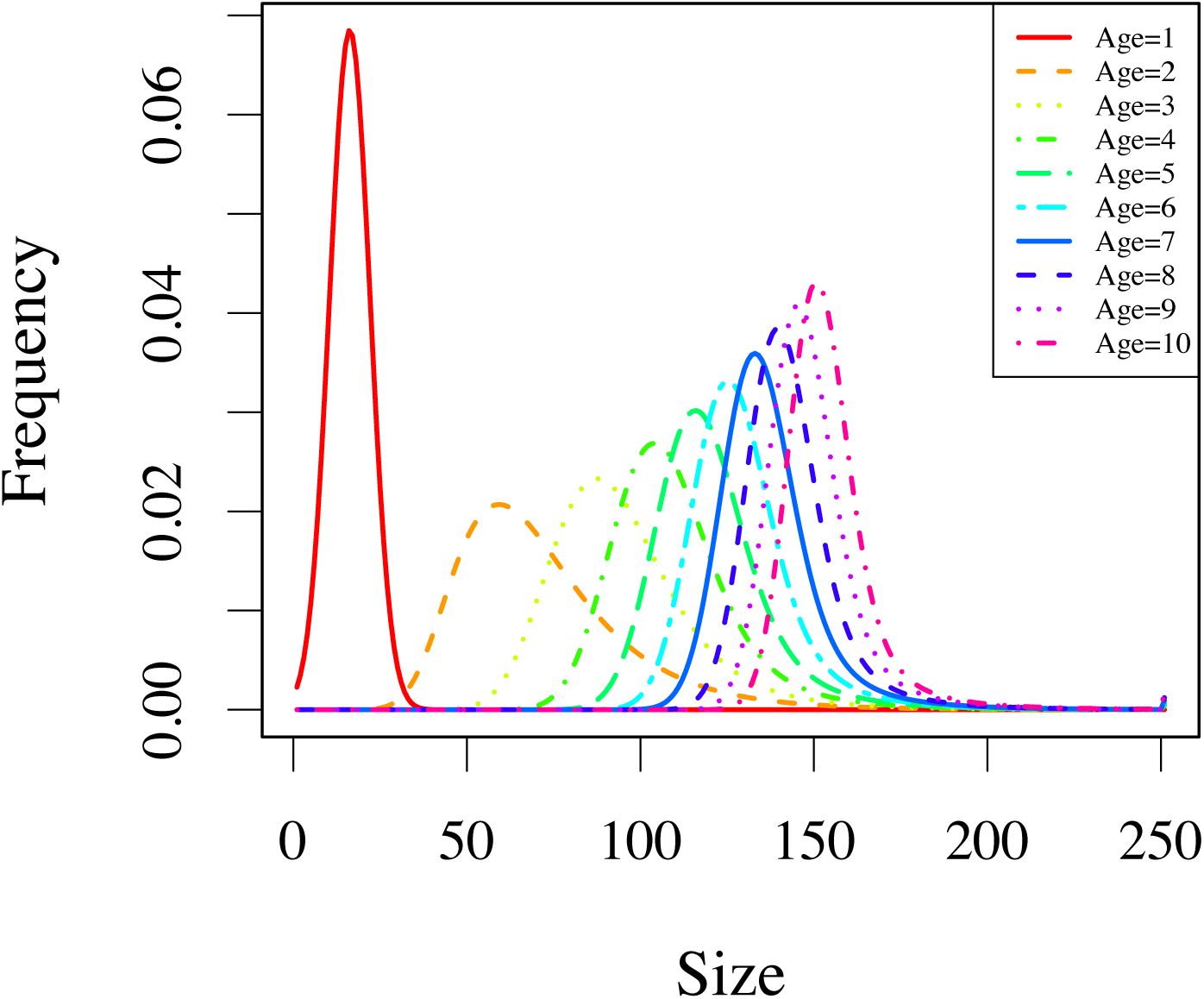
Age-specific size distributions for *C. virginica*. Size distributions for each age from *a* = 1 to *a* = 10. Distributions are generated by applying the growth and survival kernels to an initial distribution of new *a* = 1 recruits, and normalizing the distribution after each time step.

**Figure 7.**
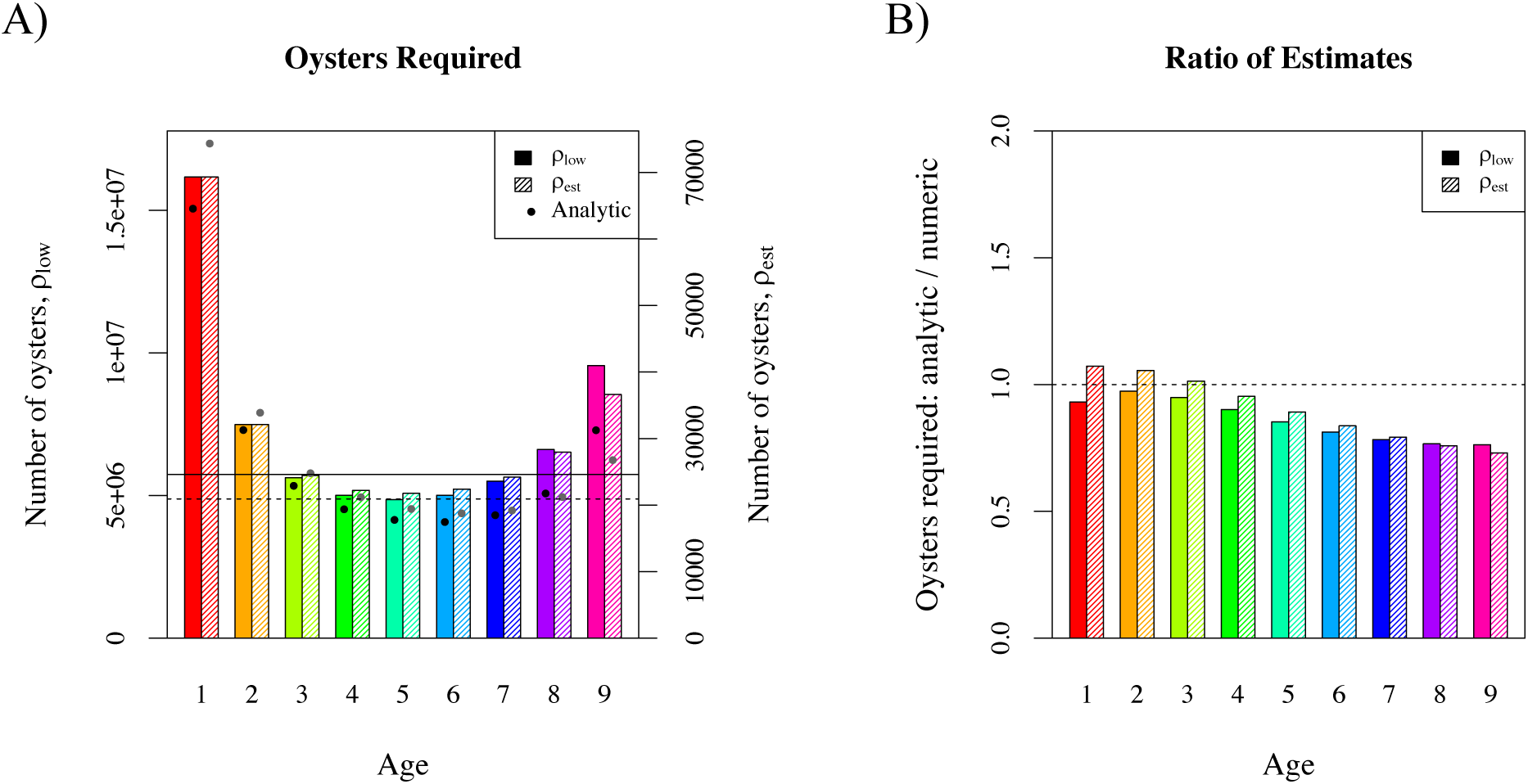
Oyster additions required for restoration. A) The required oysters of a single age, *a*, needed to push the population across the threshold surface for *ρ*_low_ (solid bars, left axis), and *ρ_est_* (shaded bars, right axis), with remaining parameters given in Table 1. Horizontal lines indicate the equilibrium total oyster number, *N̂*, for *ρ*_low_ (solid), and *ρ*_est_ (dotted). Black and gray points indicate the analytic approximation of required oysters for *ρ*_low_ and *ρ*_est_, respectively. All populations began with no available substrate, and no oysters. B) The ratio of the required oyster numbers calculated from the analytic approximation to the required oyster numbers generated from numeric simulations for each age *a* for *ρ*_low_ (solid bars, left axis), and *ρ_est_* (shaded bars, right axis). The horizontal line indicates a perfect correspondence between the two values, while bars above or below this line indicate an over- or under-estimation of the analytic approximation, respectively.

For all parameter combinations evaluated, the greatest number of oysters are required if oysters are added to the system at age 1, while the fewest number of oysters are required if oysters are added to the system between the ages of 3-7. When local retention, *ρ*, is low, significantly more oysters are required to push the population across the threshold surface, versus when *ρ* is high. Increasing *δ* increases *N̂* and thus the overall number of oysters required to cross the threshold.

While the analytic approximations work well, in general they underestimate the number of oysters required (Fig. 7B), with larger errors for older ages. When *δ* is low, the underestimation becomes more pronounced, while if *δ* is high, the analytic approximations overestimate the required oyster numbers for younger ages (particularly when *ρ* is high).

The degree of effort required for habitat enhancement depends upon the initial population size and size distribution of the population (Fig. 8). If *ρ* is low, more overall substrate is required to push the population over the threshold surface, versus when *ρ* is high. If the population is at its equilibrium size distribution, with total oyster numbers above *N̂*, the amount of substrate required is less than the equilibrium value of substrate (Fig. 8A). However, if the population begins with a size distribution similar to that of a harvested population, significantly more substrate is required beyond the equilibrium value, even if the total number of oysters begins above *N̂* (Fig. 8B).

**Figure 8.**
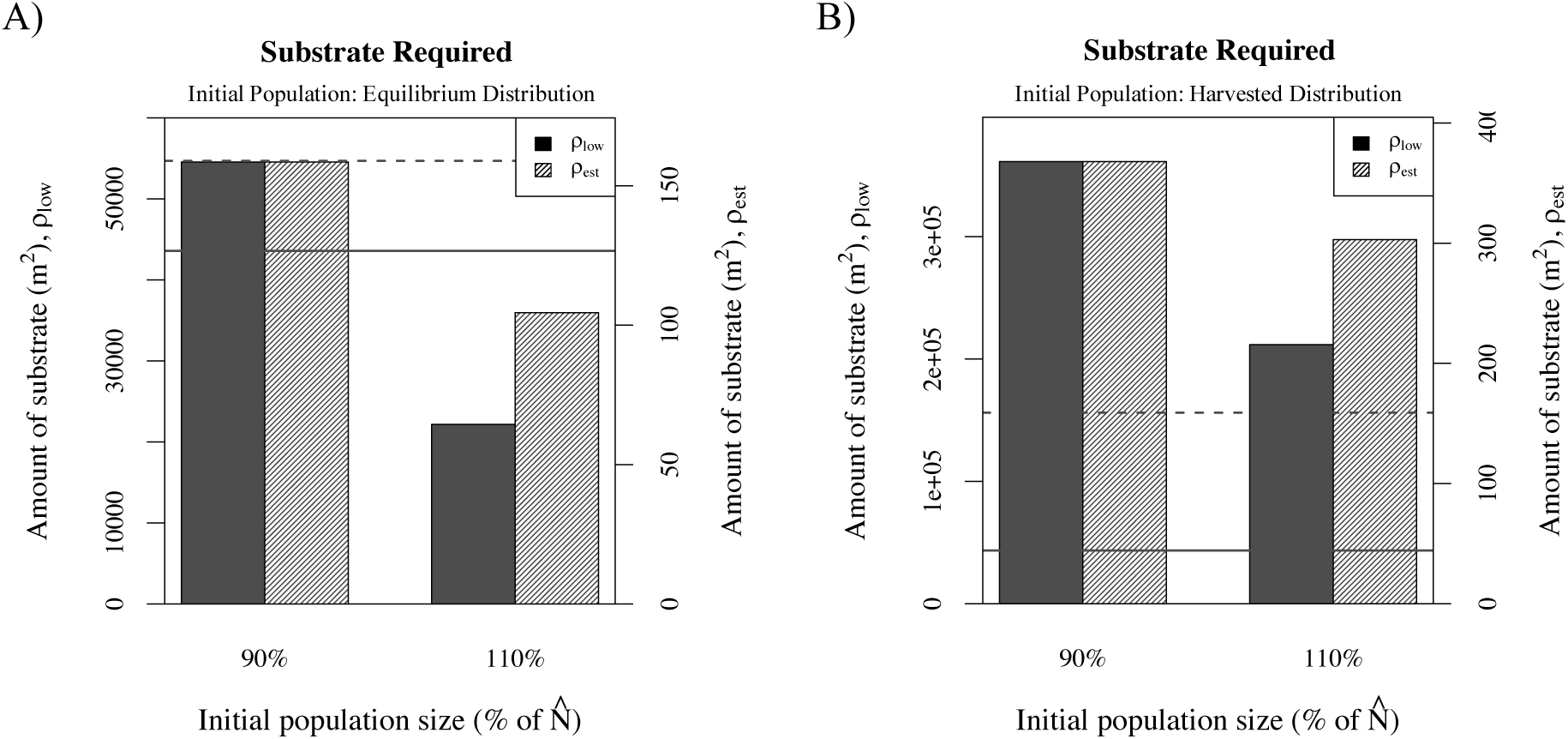
Substrate additions required for restoration. The amount of substrate required to push the population across the threshold surface for *ρ*_low_ (solid bars, left axis), and *ρ_est_* (shaded bars, right axis), with remaining parameters given in Table 1. Horizontal lines indicate the equilibrium total substrate area, *Ĥ*, for *ρ*_low_ (solid), and *ρ*_est_ (dotted). All populations began at either 90% or 110% of *N̂*, with the initial size distribution of the population set at the equilibrium size distribution (A), or the harvested size distribution (B).

## DISCUSSION

We found that incorporating positive density dependence into a size- and age-structured IPM can create an infinite dimensional threshold surface below which the population would decline to extinction, and above which the population would increase to infinity. This surface existed when *ρ* > *ρ̂*. That is, when positive feedbacks are included in the model, the maximum degree of local retention must be greater than if positive feedbacks are not considered, since the feedback term scales the number of recruits, and decreases recruitment when *Ĥ* is low.

Additionally, we found that population dynamics near the threshold surface were highly dependent upon the size and age distribution of the population. This is because the threshold not only depends on the total oyster numbers and substrate levels, but also on the size and age distribution. As such, even if the total number of oysters in the population exceeds the total equilibrium oyster number, *N̂*, the number of oysters of a particular size and age might be below the threshold surface, and the population might still decline to extinction. For example, in a population that has experienced significant harvest pressure and only oysters < 75 mm remained in the population, the total number of oysters and substrate had to be well above *N̂* and *Ĥ* in order for the population to persist. While our results indicate the existence of a restoration threshold that must be met for successful restoration (Suding and Hobbs, 2008), our results also emphasize the complexity of this threshold. It thus becomes particularly important to set a desired size- and age-structure as a goal of restoration, and not just an overall number of oysters or substrate (Baggett et al., 2014, 2015; Moore et al., 2016). In addition, in simulations that began away from the equilibrium size- and age-structure, populations exhibited oscillatory dynamics for upwards of 10 years, both when declining to extinction or persisting. When monitoring real world oyster populations, this indicates the potential difficulty of using short (approx. 5 year) time series observations to judge the need for, or the success of, restoration actions, as populations in the declining phase of an oscillation may ultimately persist.

The location of the threshold surface is dependent upon parameters whose estimates are highly uncertain. The threshold surface was most affected by changes in *ρ*, the maximum degree of local retention, with increases in *ρ* leading to decreases in the location of the threshold surface. That is, if more oyster recruits survive and remain in the natal population, the smaller the extinction region, and thus the increased likelihood of a persistent population. In terms of restoration, this supports the idea that restoration should focus on locations with a high degree of local retention or larval survival. This could be achieved by assessing the abiotic conditions of the region, such as salinity or specific hydrodynamic patterns, as well as focusing on areas of low predation and disease.

Alternatively, increases in both *δ* and *α* led to increases in the threshold surface, making it less likely that the population would persist. Increases in *δ*, the decay rate of shell substrate, indicate that the substrate will not persist as long in the system, and thus there will be less overall recruits that are able to successfully establish. This is of particular importance as there is evidence that climate change will increase ocean acidification (Orr et al., 2005; Gaylord et al., 2015). Increases in ocean acidification will decrease the calcification of oysters shells, making the shell weaker and ultimately increasing shell erosion rates (i.e. increasing *δ*; Waldbusser et al., 2011). Additionally, *δ* is temporally and spatially variable (Powell et al., 2006; Wilberg et al., 2013). Given that, restoration actions should again focus on particular locations where *δ* is low.

The *α* parameter determines the shape of the feedback function; high values of *α* decrease the strength of the positive feedback (thus increasing the equilibrium value and making it more difficult to push the population across the threshold surface), while low values of *α* increase the strength of the positive feedback. When restoring populations, there are many ways that substrate is added to existing populations. Loose shell can be dumped across large regions of the population, shells can be bagged first before being placed, or large artificial structures can be built and added to the population (Brumbaugh and Coen, 2009; Theuerkauf et al., 2015). This analysis supports the idea that the most effective technique will be the one that best facilitates oyster recruitment. While there is some data on the relationship between substrate and recruitment (Mann and Evans, 1998), much is still unknown, and oyster restoration efforts would likely benefit from additional studies investigating this relationship.

If *ρ* is low (or if *δ* is high), the location of the threshold surface will be further from the origin, and more oysters will be needed to restore the population. Additionally, our analysis shows that approximately three times as many oysters are required if spat are added to the system, rather than larger, mid-aged oysters. This might explain why seeding oysters is not always successful at enhancing oyster populations (e.g. Geraldi et al., 2013, Puckett and Eggleston, 2016). When seeding oyster populations, oyster larvae are grown on recycled shell in the lab, and then planted in the natural population once they reach a large enough size to limit mortality (Brumbaugh and Coen, 2009). As oysters grow older and larger, it becomes cost prohibitive to continue rearing the oysters in the lab. However, our results indicate that it would be worthwhile to consider methods of growing oysters to a larger size in a stress-free, high survival environment before planting them in a degraded location where restoration is desired. This could involve growing oysters off shore in protected sites for several years before moving them, or coordinating with aquaculture or community-based oyster gardening facilities.

A coupled ecological and economic modeling study conducted by Kellison and Eggleston (2004) for summer flounder stock enhancement found similar results: the number of survivors of released stock was maximized, and the total cost per survivor was minimized, when fish were released at the maximum size possible. A similar cost-benefit analysis could be done for oysters that incorporates the cost of growing a given number of oysters to a particular size to better determine the most economic size and age of oysters to use for restoration. Additionally, while this analysis looked at the minimum level of stock enhancement required for persistence, future work could also incorporate an “economic restoration threshold” (Lampert and Hastings, 2014) to determine the optimal level of stock enhancement (which might exceed the minimum level) required to meet a restoration goal in a cost-effective manner.

For substrate addition, an unrealistic amount of substrate (> 13 million m^2^) was required if the population began at low (<10% of *N̂*) population levels (results not shown). If the population began close to the equilibrium total oyster numbers, the amount of substrate required was closer to the equilibrium substate levels. However, if *ρ* was low or *δ* was high, substantially more substrate was required. The initial size distribution of the population was also important. If the population began at the equilibrium size distribution, then the amount of substrate required was less than the equilibrium substrate level if the population began at 110% of *N̂*, while the amount was greater than or equal to the equilibrium substrate level if the population began at 90% of *N̂*. However, if the population began with a size distribution similar to that of a harvested population, even if the population began above the equilibrium number of total oysters, significantly more substrate beyond the equilibrium substrate levels was required to restore the population. This result reinforces the idea that the structure of the population is equally important as the overall size of the population for a healthy, persistent population (Baggett et al., 2014, 2015; Moore et al., 2016).

The analytic approximation of oysters required tended to underestimate the number of oysters required, particularly for older ages and when *δ* is low. However, within the range of parameter values explored, in most cases the analytic approximation was within 25% of the numeric value, with the worst case still within 40% of the numeric value. Given the high dimension of the threshold surface, it is surprising that the analytic approximation performs this well. This success of the analytical approach suggests that it might be useful for other IPMs and matrix models with positive feedbacks.

### Limitations and challenges

Our model, which includes both structuring population variables and positive density dependence, allows for direct assessment of the required number of oysters or substrate for a persistent population. However, there are several important factors that are not yet incorporated into the model presented here. First, we only include positive feedbacks in the fecundity term of the model. In reality, the amount of shell substrate will also have a positive effect on adult oyster growth and survival, for example, through the interaction between reef height and shape, water depth, and water flow speeds (Lenihan and Peterson, 1998; Lenihan, 1999; Bartol et al., 1999), though it’s likely these effects will be small relative to the positive density dependence effect on recruitment. Additionally, the three-dimensional shape of a reef plays a role in determining how much of the overall shell surface area is available for settlement. Future work could extend the IPM presented here to include a variable for reef height or shape, or a structuring variable that represents the location of individual oysters within the reef. This will likely significantly increase the complexity of the dynamics.

Our model also does not include negative density dependence, which is important for oyster dynamics (Knights and Walters, 2010; Puckett and Eggleston, 2012). However, preliminary analysis of a model with both positive and negative feedbacks on fecundity indicate that, with the exception of a positive stable equilibrium surface in addition to the unstable threshold equilibrium surface, qualitative results are similar. Additionally, since we are focused on restoring highly degraded populations, population sizes are likely too small for negative density dependence to have a large effect.

Next, our model assumes a closed population with no external subsidy of recruits. Because of this, any new oysters must either be generated by the local population, or added through restoration actions. This likely explains the unrealistic amount of substrate required at low population sizes. Though many natural oyster reefs are fairly isolated, such as the site used to parameterize the model, many natural oyster reefs receive a large proportion of recruits from external populations, and even isolated populations likely receive some larvae from external sources (Lipcius et al., 2008, 2011; Puckett and Eggleston, 2016). Future work could extend the model to allow for external recruitment to better understand how external subsidies affect the location of the threshold surface. Additionally, the inclusion of external recruitment into future models can give managers a better sense for the relative effectiveness of either stock or habitat enhancement.

Finally, model parameters *ρ*, *δ*, and *α* are highly uncertain, and also highly variable in space and time. While qualitative results do not differ significantly across the range of parameter values explored, if managers are interested in determining more precisely the location of the threshold surface, more accurate parameter estimates are needed. In most cases, managers will not have a firm grasp on any of the three parameters, but based on our elasticity analysis, obtaining accurate estimates of local retention should be prioritized followed by the relationship between substrate and recruitment, and the local substrate decay rates. Additionally, to incorporate variability in model parameters, future work could extend the model to allow for stochasticity, particularly in fecundity and recruitment, which is highly variable both within and between years (Cox and Mann, 1992; Ortega and Sutherland, 1992; Siegel et al., 2008; Mroch et al., 2012). Lastly, the model implements a restoration strategy in year 1 of a 150 year time line. Future work could investigate restoration actions over multiple years, as well simultaneous substrate and oyster addition.

### Conclusion

Using demographic data from a population of Eastern oyster, *C. virginica*, in Pamlico Sound, North Carolina, our modeling analysis indicates the importance of positive density dependence at influencing population dynamics. We show how population parameters, such as local retention and the decay rate of shell substrate, influence the amount of restoration needed to restore a degraded population. We also find that if mid-aged oysters are used for stock enhancement of fully degraded populations, fewer numbers are required for restoration than if oyster spat are used. Finally, we find that restoration of existing populations depends strongly upon the initial size distribution of the population. Future work allowing for external recruitment is needed to better investigate the relative importance of stock enhancement versus habitat enhancement.

## ACKNOWLEDGMENTS

Support for J.L. Moore and S.J. Schreiber was provided by NSF Grants DMS-1022639 and DMS-1313418. Support for B. Puckett was provided by NSF award OCE-1155628 and NC Sea Grant award 14-HCE-9.

## DESCRIPTION OF SUPPORTING INFORMATION

Appendix S1. **Details of elasticity analysis.** Provides explicit equations used in elasticity analysis for *ρ*, *δ*, and *α*.

Appendix S2. **Characteristics of the threshold surface at the equilibrium.** Shows that the tangent space of the threshold surface at the equilibrium, **x̂**, is perpendicular to the left, dominant eigenvalue of the derivative operator evaluated at **x̂**.

Appendix S3. **Analytic approximation of required oysters.** Provides a discussion of how the required number of age-specific oysters was approximated analytically.

## Appendix S1. Details of Elasticity Analysis.

In this Appendix, we discuss the specific equations used to determine the elasticity of the threshold surface to *ρ*, *δ*, and *α*.

Let **x** = (**n**, *H*)^*T*^ and **n** = (*n*_1_(*y*), *B*_1_, ⋯, *n_A_*(*y*), *B_A_*), where

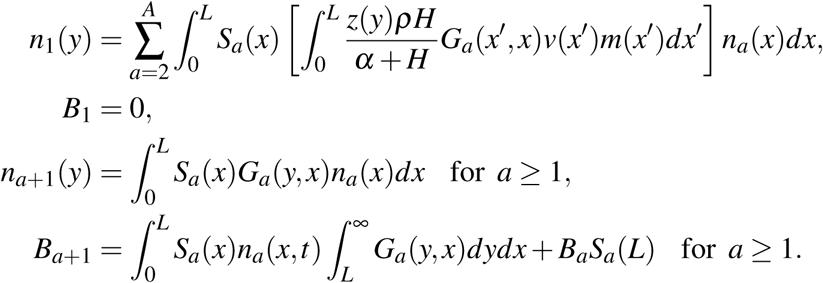

Then,

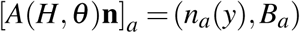

and

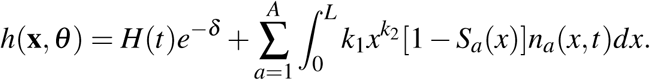

As described in the text, the sensitivity of **x̂** = (**n̂**, *Ĥ*) to *θ* is

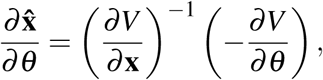

where

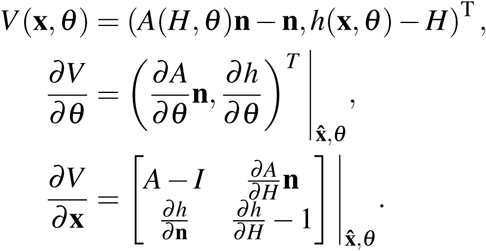

Let

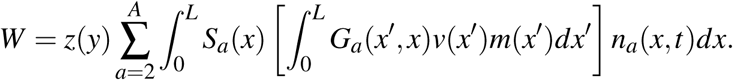

Then,

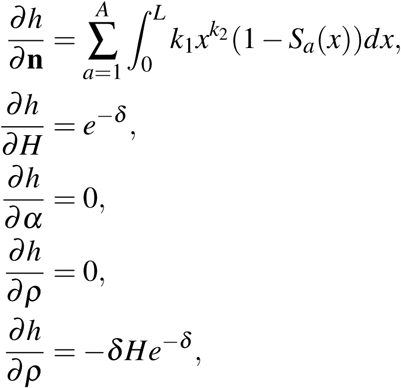

and,

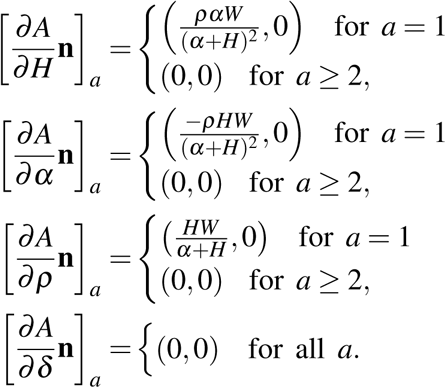

Elasticities are then given by

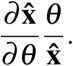

## APPENDIX S2. Characteristics of Threshold Surface.

In this Appendix, we show that the tangent space of the threshold surface at the equilibrium **x̂** is perpendicular to the left, dominant eigenvalue of the derivative operator evaluated at **x̂**. Recall that **x** = (**n**, *H*) and the dynamics of the IPM is given by iterating the map *F*(**x**) = (*A*(*H*)**n**, *h*(**x**)). The Jacobian matrix of *F* at **x̂** is given by

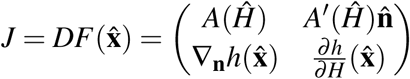

where ∇_**n**_*h* denotes the gradient of *h* with respect **n**. This operator *J* is non-negative and power positive as *A′* (*H*) is strictly positive in the age 1 class and 0 elsewhere, ∇_**n**_*h* is strictly positive and 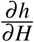 is strictly positive. Let *λ* be the dominant eigenvalue of *J* and **v** and **w** be corresponding left and right eigenvectors.

To show that tangent space of the threshold surface at **x̂** is perpendicular to **v**, it suffices to show for any right eigenvector, call it **u**, not spanned by **w** is perpendicular to **w**. Let *μ* ≠ *λ* be the eigenvalue associated with **u**. Then

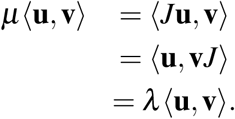

As *λ* ≠ *μ*, it follows that 〈**u**, **v**〉 = 0.

## Appendix S3. Analytic Approximation of Required Oyster Additions.

In this Appendix, we derive the analytic approximation of the number of age-specific oysters required to cross the threshold surface, beginning with no oysters, and no substrate. Let **x̂** = (**n̂**, *Ĥ*) be the vector representing the equilibrium, **w** be the left dominant eigenvector that is perpendicular to the linearized threshold surface, and **v_a_** be the vector given by the normalized size distribution of oysters of age *a*. We are then interested in determining *b_a_*, the multiplier of **v_a_** such that ∥*b_a_***v_a_**∥ just crosses the linearized threshold surface (Fig. S1). We solve for *b_a_* by first determining angles *B* and *C*, where

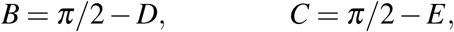

and

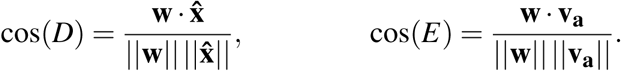

Then, using the Law of Sines,

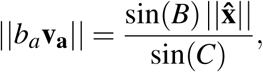

and thus

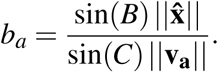

We then calculate *b_a_* for each age from *a* = 1 to *a* = 9.

**Figure S1.**
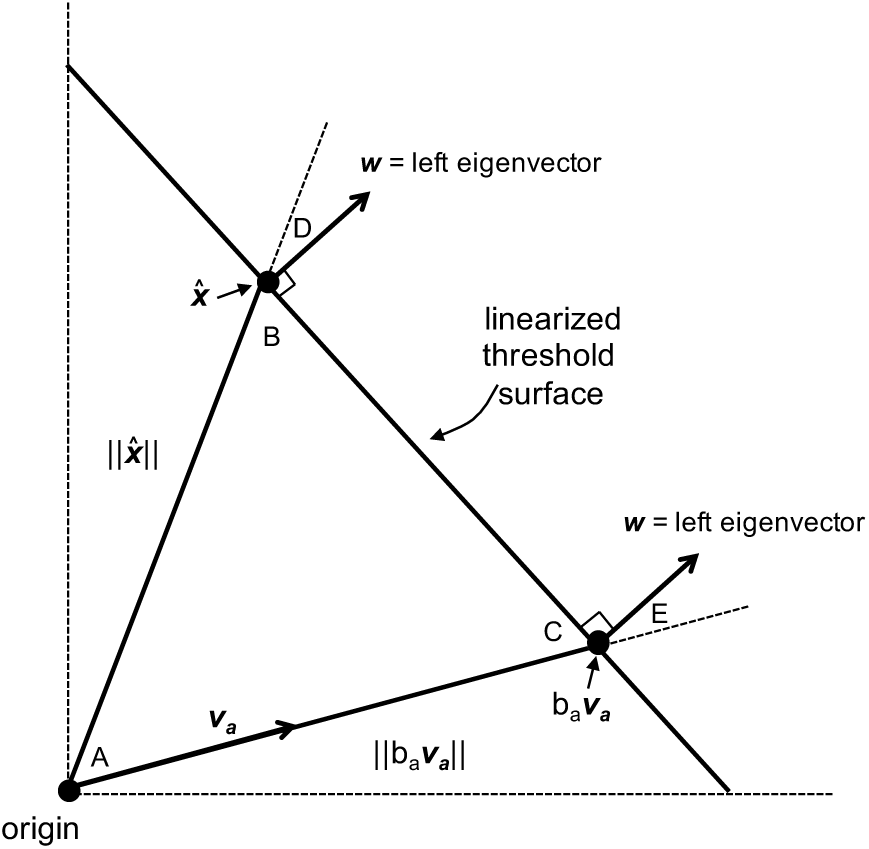

## Literature Cited

L Airoldi and MW Beck. Loss, status and trends for coastal marine habitats of Europe. Oceanography and Marine Biology: An Annual Review, 45:345–405, 2007.

WC Allee. Principles of animal ecology. University of Chicago Press, 1931.

WC Allee. Animal aggregations: a study in general sociology. Saunders, 1949.

LP Baggett, SP Powers, R Brumbaugh, LD Coen, B DeAngelis, J Greene, B Hancock, and S Morlock. Oyster habitat restoration monitoring and assessment handbook. The Nature Conservancy, 2014.

LP Baggett, SP Powers, RD Brumbaugh, LD Coen, BM DeAngelis, JK Greene, BT Hancock, SM Morlock, BL Allen, DL Breitburg, D Bushek, JH Grabowski, RE Grizzle, ED Grosholz, MK La Peyre, MW Luckenbach, KA McGraw, MF Piehler, SR Westby, and PSE zu Ermgassen. Guidelines for evaluating performance of oyster habitat restoration. Restoration Ecology, 23(6):737–745, 2015.

IK Bartol, R Mann, and M Luckenbach. Growth and mortality of oysters (*Crassostrea virginica*) on constructed intertidal reefs: effects of tidal height and substrate level. Journal of Experimental Marine Biology and Ecology, 237(2):157–184, 1999.

MW Beck, RD Brumbaugh, L Airoldi, A Carranza, LD Coen, C Crawford, O Defeo, GJ Edgar, B Hancock, MC Kay, S Hunter, MW Luckenbach, CL Toropova, and GG Zhang. Oyster reefs at risk and recommendations for conservation, restoration, and management. BioScience, 61 (2):107–116, 2011.

BE Beisner, DT Haydon, and K Cuddington. Alternative stable states in ecology. Frontiers in Ecology and the Environment, 1(7):376–382, 2003.

RD Brumbaugh and LD Coen. Contemporary approaches for small-scale oyster reef restoration to address substrate versus recruitment limitation: a review and comments relevant for the Olympia oyster, *Ostrea lurida* Carpenter 1864. Journal of Shellfish Research, 28(1):147–161, 2009.

JE Byers, K Cuddington, CG Jones, TS Talley, A Hastings, JG Lambrinos, JA Crooks, and WG Wilson. Using ecosystem engineers to restore ecological systems. Trends in Ecology and Evolution, 21(9):493–500, 2006.

RB Carnegie and EM Burreson. Declining impact of an introduced pathogen: *Haplosporidium nelsoni* in the oyster *Crassostrea virginica* in Chesapeake Bay. Marine Ecology Progress Series, 432:1–15, 2011.

LD Coen, RD Brumbaugh, D Bushek, R Grizzle, MW Luckenbach, MH Posey, SP Powers, and SG Tolley. Ecosystem services related to oyster restoration. Marine Ecology Progress Series, 341:303–307, 2007.

F Courchamp, T Clutton-Brock, and B Grenfell. Inverse density dependence and the Allee effect. Trends in Ecology and Evolution, 14(10):405–410, 1999.

F Courchamp, L Berec, and J Gascoigne. Allee Effects in Ecology and Conservation. Oxford University Press, 2008.

C Cox and R Mann. Temporal and spatial changes in fecundity of eastern oysters, *Crassostrea virginica* (Gmelin, 1791) in the James River, Virginia. Journal of Shellfish Research, 11(1): 49–54, 1992.

K Cuddington, WG Wilson, and A Hastings. Ecosystem engineers: feedback and population dynamics. The American Naturalist, 173(4):488–498, 2009.

PS Galtsoff. *The American oyster* Crassostrea virginica *Gmelin*. Fishery Bulletin of the Fish and Wildlife Service, vol. 64, 1964.

PS Galtsoff. The American Oyster, *Crassostrea virginica* Gmelin – morphology and structure of shell. Fishery Bulletin, 64:16–47, 1966.

J Gascoigne and RN Lipcius. Allee effects in marine systems. Marine Ecology Progress Series, 269:49–59, 2004.

B Gaylord, KJ Kroeker, JM Sunday, KM Anderson, JP Barry, NE Brown, SD Connell, S Dupont, et al. Ocean acidification through the lens of ecological theory. Ecology, 96(1):3–15, 2015.

NR Geraldi, M Simpson, SR Fegley, P Holmlund, and CH Peterson. Addition of juvenile oysters fails to enhance oyster reef development in Pamlico Sound. Marine Ecology Progress Series, 480:119–129, 2013.

JH Grabowski, RD Brumbaugh, RF Conrad, G Andrew, JJ Opaluch, CH Peterson, MF Piehler, SP Powers, and AR Smyth. Economic valuation of ecosystem services provided by oyster reefs. BioScience, 62(10):900–909, 2012.

A Hastings and DB Wysham. Regime shifts in ecological systems can occur with no warning. Ecology Letters, 13(4):464–72, 2010.

EE Hofmann, D Bushek, S Ford, X Guo, D Haidvogel, D Hedgecock, JM Klinck, C Milbury, D Narvaez, EN Powell, Y Wang, Z Wang, J Wilkin, and L Zhang. Understanding how disease and environment combine to structure resistance in estuarine bivalve populations. Oceanography, 22(4):212–231, 2009.

RM Housego and JH Rosman. A model for understanding the effects of sediment dynamics on oyster reef development. Estuaries and Coasts, 39(2):495–509, 2016.

WC Jordan-Cooley, RN Lipcius, LB Shaw, J Shen, and J Shi. Bistability in a differential equation model of oyster reef height and sediment accumulation. Journal of Theoretical Biology, 289: 1–11, 2011.

GT Kellison and DB Eggleston. Coupling ecology and economy: modeling optimal release scenarios for summer flounder (*Paralichthys dentatus*) stock enhancement. Fishery Bulletin, 102:78–93, 2004.

VS Kennedy, DL Breitburg, MC Christman, MW Luckenbach, K Paynter, J Kramer, KG Sellner, J Dew-Baxter, C Keller, and R Mann. Lessons learned from efforts to restore oyster population in Virginia and Maryland. Journal of Shellfish Research, 30:1–13, 2011.

AM Knights and K Walters. Recruit-recruit interactions, density-dependent processes and population persistence in the Eastern oyster *Crassostrea virginica*. Marine Ecology Progress Series, 404:79–90, 2010.

A Lampert and A Hastings. Optimal control of population recovery - the role of economic restoration threshold. Ecology Letters, 17:28–35, 2014.

HS Lenihan. Physical-biological coupling on oyster reefs: how habitat structure influences individual performance. Ecological Monographs, 69(3):251–275, 1999.

HS Lenihan and CH Peterson. How habitat degredation through fishery disturbance enhances impacts of hypoxia on oyster reefs. Ecological Applications, 8(1):128–140, 1998.

HS Lenihan, CH Peterson, and JM Allen. Does flow speed also have a direct effect on growth of active suspension-feeders: an experimental test on oysters. Limnology and Oceanography, 41 (6):1359–1366, 1996.

RN Lipcius, DB Eggleston, SJ Schreiber, RD Seitz, J Shen, M Sisson, WT Stockhausen, and HV Wang. Importance of metapopulation connectivity to restocking and restoration of marine species. Reviews in Fisheries Science, 16(1-3):101–110, 2008.

RN Lipcius, GM Ralph, J Liu, V Hill, AT Morzillo, JA Wiens, et al. Evidence of source-sink dynamics in marine and estuarine species. Sources, sinks and sustainability, pages 361–381, 2011.

RN Lipcius, RP Burke, DN McCulloch, SJ Schreiber, DM Schulte, RD Seitz, and J Shen. Overcoming restoration paradigms: value of the historical record and metapopulation dynamics in native oyster restoration. Frontiers in Marine Science, 2:65, 2015.

R Mann and DA Evans. Estimation of oyster, *Crassostrea virginica*, standing stock, larval production and advective loss in relation to observed recruitment in the James River, Virginia. Journal of Shellfish Research, 17(1):239–253, 1998.

R Mann and EN Powell. Why oyster restoration goals in the Chesapeake Bay are not and probably cannot be achieved. Journal of Shellfish Research, 26(4):905–917, 2007.

JL Moore, RN Lipcius, B Puckett, and SJ Schreiber. The demographic consequences of growing older and bigger in oyster populations. Ecological Applications, 26(7):2206–2217, 2016.

RM Mroch, DB Eggleston, and BJ Puckett. Spatiotemporal variation in oyster fecundity and reproductive output in a network of no-take reserves. Journal of Shellfish Research, 31(4): 1091–1101, 2012.

JA Nestlerode, MW Luckenbach, and FX O’Beirn. Settlement and survival of the oyster *Crassostrea virginica* on created oyster reef habitats in Chesapeake Bay. Restoration Ecology, 15(2):273–283, 2007.

M Nystrom, AV Norstrom, T Blenckner, M de la Torre Castro, JS Eklof, C Folke, H Osterblom, RS Steneck, M Thyresson, and M Troell. Confronting feedbacks of degraded marine ecosystems. Ecosystems, 15:695–710, 2012.

JC Orr, VJ Fabry, O Aumont, L Bopp, SC Doney, RA Feely, A Gnanadesikan, Gruber N, et al. Anthropogenic ocean acidification over the twenty-first century and its impact on calcifying organisms. Nature, 437(29):681–686, 2005.

S Ortega and JP Sutherland. Recruitment and growth of the eastern oyster *Crassostrea virginica*, in North Carolina. Estuaries, 15(2):158–170, 1992.

EN Powell, JN Kraeuter, and KA Ashton-Alcox. How long does oyster shell last on an oyster reef? Estuarine, Coastal and Shelf Science, 69:531–542, 2006.

EN Powell, JM Klinck, KA Ashton-Alcox, and JN Kraeuter. Multiple stable reference points in oyster populations: implications for reference point-based management. Fishery Bulletin, 107 (2):133–147, 2009a.

EN Powell, JM Klinck, KA Ashton-Alcox, and JN Kraeuter. Multiple stable reference points in oyster populations: biological relationships for the eastern oyster (*Crassostrea virginica*) in Delaware Bay. Fishery Bulletin, 107(2):109–132, 2009b.

SP Powers, CH Peterson, JH Grabowski, HS Lenihan, et al. Success of constructed oyster reefs in no-harvest sanctuaries: implications for restoration. Marine Ecology Progress Series, 389: 159–170, 2009.

BJ Puckett and DB Eggleston. Oyster demographics in a network of no-take reserves: recruitment, growth, survival, and density dependence. Marine and Coastal Fisheries, 4(1): 605–627, 2012.

BJ Puckett and DB Eggleston. Metapopulation dynamics guide marine reserve design: importance of connectivity, demographics, and stock enhancement. Ecosphere, 7(6):e01322, 2016.

BJ Puckett, DB Eggleston, PC Kerr, and RA Luettich Jr. Larval dispersal and population connectivity among a network of marine reserves. Fisheries Oceanography, 23(4):342–361, 2014.

R Core Team. R: A Language and Environment for Statistical Computing. R Foundation for Statistical Computing, Vienna, Austria, 2015.URL https://www.R-project.org/.

BJ Rothschild, JS Ault, P Goulletquer, and M Heral. Decline of the Chesapeake Bay oyster population: a century of habitat destruction and overfishing. Marine Ecology Progress Series, 111:29–39, 1994.

M Scheffer and SR. Carpenter. Catastrophic regime shifts in ecosystems: linking theory to observation. Trends in Ecology & Evolution, 18(12):648–656, 2003.

M Scheffer, S Carpenter, JA Foley, C Folke, and B Walker. Catastrophic shifts in ecosystems. Nature, 413(6856):591–6, 2001.

SJ Schreiber. Allee effects, extinctions, and chaotic transients in simple population models. Theoretical Population Biology, 64:201–209, 2003.

SJ Schreiber. On Allee effects in structured populations. Proceedings of the American Mathematical Society, 132(10):3047–3053, 2004.

DM Schulte, RP Burke, and RN Lipcius. Unprecedented restoration of a native oyster metapopulation. Science, 325:1124–1128, 2009.

DA Siegel, S Mitarai, CJ Costello, SD Gaines, BE Kendall, RR Warner, and KB Winters. The stochastic nature of larval connectivity among nearshore marine populations. PNAS, 105(26): 8974–8979, 2008.

PA Stephens, WJ Sutherland, and RP Freckleton. What is the Allee effect? Oikos, 87(1): 185–190, 1999.

KN Suding and RJ Hobbs. Threshold models in restoration and conservation: a developing framework. Trends in Ecology and Evolution, 24(5):271–279, 2008.

J Taylor and D Bushek. Intertidal oyster reefs can persist and function in a temperate North American Atlantic estuary. Marine Ecology Progress Series, 361:301, 2008.

SJ Theuerkauf, RP Burke, and RN Lipcius. Settlement, growth, and survival of eastern oysters on alternative reef substrates. Journal of Shellfish Research, 34(2):241–250, 2015.

GG Waldbusser, RA Steenson, and MA Green. Oyster shell dissolution rates in estuarine waters: effects of pH and shell legacy. Journal of Shellfish Research, 30(3):659–669, 2011.

MJ Wilberg, JR Wiedenmann, and JM Robinson. Sustainable exploitation and management of autogenic ecosystem engineers: application to oysters in Chesapeake Bay. Ecological Applications, 23(4):766–776, 2013.

JL Williams, TEX Miller, and SP Ellner. Avoiding unintentional eviction from integral projection models. Ecology, 93(9):2008–2014, 2012.

PSE zu Ermgassen, MD Spalding, B Blake, LD Coen, B Dumbauld, S Geiger, JH Grabowski, R Grizzle, M Luckenbach, K McGraw, W Rodney, JL Ruesink, SP Powers, and R Brumbaugh. Historical ecology with real numbers: past and present extent and biomass of an imperilled estuarine habitat. Proceedings of the Royal Society B: Biological Sciences, 279(1742): 3393–3400, 2012.

